# The effect of overnight consolidation in the perceptual learning of non-native tonal contrasts

**DOI:** 10.1101/731042

**Authors:** Zhen Qin, Caicai Zhang

**Affiliations:** Department of Chinese and Bilingual Studies, The Hong Kong Polytechnic University, Hong Kong SAR, China; Research Center for Language, Cognition, and Neuroscience, The Hong Kong Polytechnic University, Hong Kong SAR, China

## Abstract

Sleep-mediated overnight consolidation was found to facilitate perceptual learning by promoting learners’ generalization across talkers in their perception of novel segmental categories. Lexical tone differs from most segmental contrasts in that it is highly variable across talkers, and displays dynamic change over time. It remains unclear whether a similar (or a larger) effect of overnight consolidation would be found for perceptual learning of novel tonal contrasts. Thus, this study aims to examine whether overnight consolidation facilitates generalization across talkers in the discrimination and identification of novel Cantonese level tones by Mandarin listeners. Two groups of Mandarin listeners were perceptually trained either in the morning or in the evening. Listeners were trained in a tone identification (ID) task using stimuli produced by a trained talker. Their development was then tested in the ID and AX discrimination tasks using stimuli produced by trained and untrained talkers in three posttests following training: immediately after training, 12-hour delay, and 24-hour delay. While the evening group slept between the first and second posttests, the morning group did not. The results of accuracy rates in the ID task showed that while Mandarin listeners trained in the evening showed an improved trend, predicted by their individual sleep time, in identifying the level tones produced by both the trained and untrained talkers, Mandarin listeners trained in the morning showed a declining trend. In contrast, the results of d-prime scores in the AX discrimination task did not show different developmental patterns between the two groups. Consistent with previous studies on segmental learning, the finding suggests that overnight consolidation might have assisted the evening trainees’ formation of a more abstract (talker-independent) representation of novel tone categories in memory traces. The results are discussed regarding the features of lexical tones to shed light on the mechanism of phonetic learning.

## 1 Introduction

Converging evidence indicates that sleep supports various aspects of language learning by facilitating the memory consolidation of newly learned knowledge [see 1,2 for review]. Sleep-mediated memory consolidation (i.e., overnight consolidation) plays an important role in novel word learning by facilitating the learning process through an overnight interval [3–7]. For instance, one study found that only the set of novel words learned in the evening and consolidated during the overnight interval, compared with another set of novel words learned during the daytime without overnight consolidation, showed a lexical competition with existing words and elicited faster responses than those learned during the daytime in the set of assessment tasks [6].

Non-native speech sounds are perceptually difficult for adults to learn, particularly when the sounds are similar, but not identical, to the sounds of their native language (L1) [8,9]. However, the conditions under which the learning of new sounds is facilitated or inhibited are unclear. A growing literature showed that overnight consolidation facilitates listeners’ perceptual learning of non-native sound categories [10–14]. In several studies, Earle and her colleagues examined how learned sound categories were encoded into long-term memory following a session of laboratory training by focusing on the effect of overnight consolidation in the identification and discrimination of novel phonetic categories, that is, a Hindi dental and retroflex stop contrast [15–19]. Specifically, Earle and Myers showed that overnight consolidation promotes English listeners’ generalization across talkers in their identification, but not in the discrimination, of the novel Hindi contrast [16]. A pretest-training-posttest paradigm was conducted on two groups of English listeners who were perceptually trained either in the morning or in the evening, following this procedure: (i) listeners were perceptually trained using stimuli produced by a trained talker in an identification task; (ii) listeners’ perceptual development was tested in the identification and AX discrimination tasks using stimuli produced by trained and untrained talkers. Listeners’ perceptual development were assessed in series of posttests at three time points over 24h following training: immediately after training, 12-hour delay, and 24-hour delay. Whereas the evening group slept between the first and second posttests, the morning group did not. The results of identification tests showed that while the English listeners who were trained in the evening improved significantly their performance in identifying the stop stimuli produced by the untrained talker, those trained in the morning did not show such a pattern. In contrast, the two training groups did not show perceptual changes in identifying the stop stimuli produced by the trained talker. The results of the AX discrimination tests did not show a difference of the two training groups in their perceptual development. The discrepancy between the identification and discrimination tests was attributed to the greater variability (untrained vowel contexts) of the discrimination tests, which may have made it difficult for listeners to use consistent criteria in performing the tasks. The findings of the identification tests indicate that the overnight consolidation process should have facilitated the abstraction of novel sound categories, that is, a transfer of novel sound information from an acoustic-sensory-based trace to a more abstract (i.e., talker-independent) representation of the target stop contrast.

A similar effect of overnight consolidation in the abstraction of novel phonetic information was also found in lexically-guided phonetic retuning of non-native segments. For example, Xie, Earle and Myers conducted a perceptual learning paradigm (an exposure phase and a test phase) on English listeners who received exposure to English words ending in /d/ (e.g., seed; no words ending in /t/) [20]. Participants received the input either in the morning (Same-Day group) or in the evening (Overnight group) during an exposure phase, and then were assessed in a /d/-/t/ (e.g., seed vs. seat) identification task 12 hours later as a test phase. While the stimuli from a trained Mandarin accented talker was used during the exposure phase, the stimuli from both the trained talker and an untrained Mandarin accented talker who are not phonetically similar were used in the identification task of the test phase. The results of the identification task showed that the Overnight group showed improvements in terms of the percentage of /d/ responses for the untrained talker 12 hours later. In contrast, the Same-Day group’s performance in the percentage of /d/ responses declined with a 12-hour delay. In line with the results of previous studies [16], the findings would suggest that overnight consolidation facilitated generalization of accent adaptation to a new (i.e., untrained) Mandarin talker by helping English listeners abstract away from specific properties of the trained talker in a lexically-guided phonetic retuning process.

The overnight consolidation effect in talker generalization (i.e., abstraction) found for the perceptual learning and lexically-guided phonetic retuning of non-native segments, albeit informative, raise the question of whether a similar (or a larger) effect of overnight consolidation in talker generalization would be found in a perpetual learning of non-native tones. Different from segments (consonants and vowels), as illustrated in Fig 1, lexical tones are highly dynamic, requiring listeners to evaluate the pitch they hear in the signal against the pitch range of the talker and continuously update this evaluation as more of the pitch contour is heard over time [21–25]. Importantly, a given talker shows variability in the production of tonal categories, thus also requiring listeners to evaluate the pitch they hear against the talker’s different realizations of the same tonal categories (within-category variations) and of different tonal categories (cross-category variations) [26–32]. Thus, listeners need to extract the abstract representations from tonal exemplars in order to generalize successfully across talkers [27]. The variable and dynamic features of lexical tones make it worth investigating whether listeners who learn novel lexical tones with overnight consolidation would show a facilitated generalization across talkers at least to a similar degree as that reported on the learning of novel segmental contrasts.

**Fig 1.**
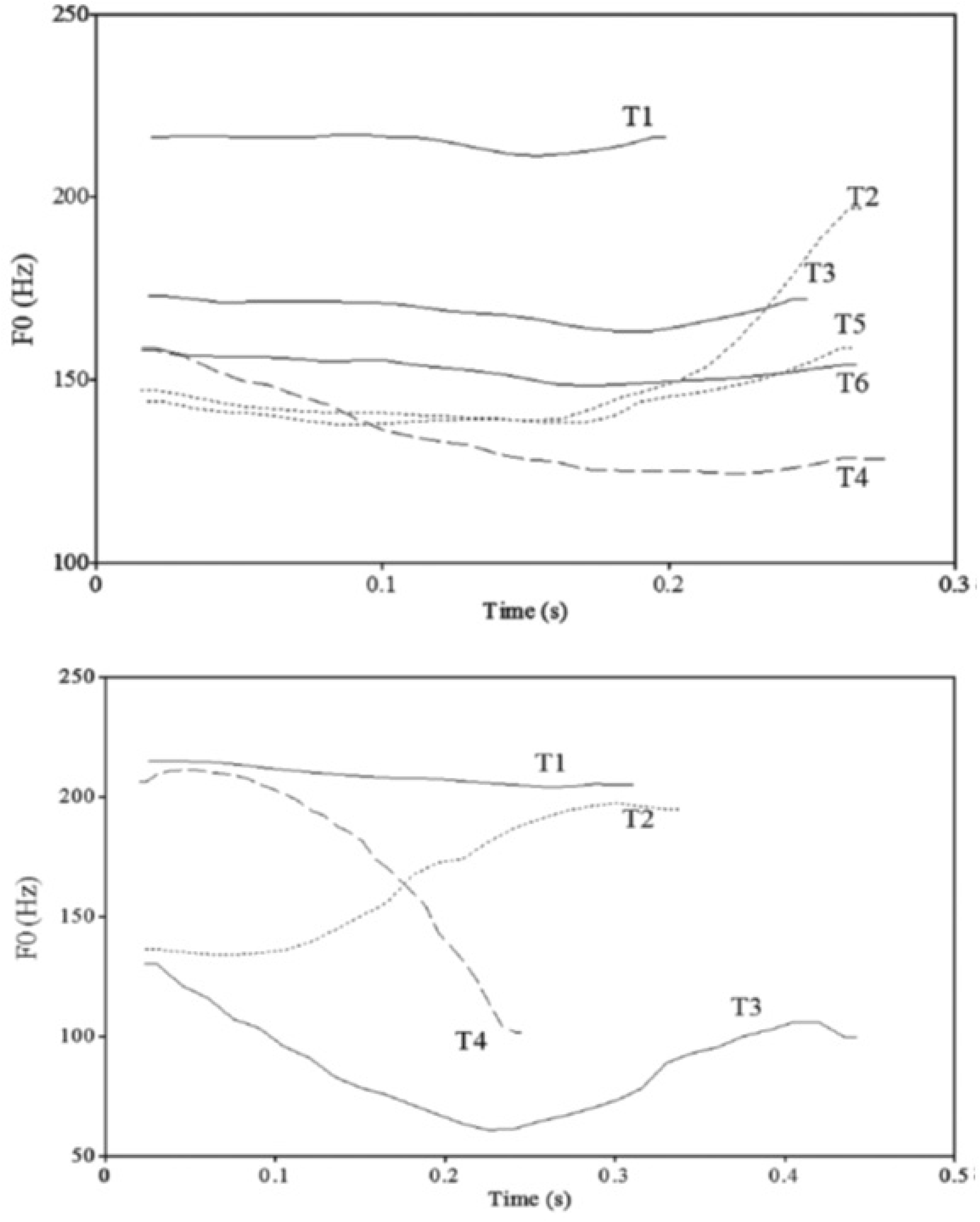
Tone contours of the six Cantonese tones and four Mandarin tones. The Cantonese tones (top panel) were produced by a female native speaker of Hong Kong Cantonese. The Mandarin tones (bottom panel) were produced by a male native speaker of Beijing Mandarin. Both figures were adopted from Qin & Jongman, 2016 for the purpose of illustration.

Suprasegmental information (e.g., pitch) is important for distinguishing among words in Chinese languages. For instance, as illustrated in Fig 1, segmentally identical words that contain different lexical tones in Cantonese (/si 55/ ‘silk (Tone 1 (T1)), /si 25/ ‘history’(T2), /si 33/ ‘to try’(T3), /si 21/ ‘time (T4), /si 23/ ‘city’(T5), /si 22/ ‘matter’(T6) [33]) and in Mandarin (e.g., /pā/ ‘eight’ (T1), /pá/ ‘to pull out’ (T2), /pǎ/ ‘to hold’ (T3), /pà/ ‘father’ (T4) [34]) differ in meaning. While pitch is the primary cue in tone perception, listeners with different language backgrounds attend to different pitch dimensions under the influence of their L1 prosodic system [35–38,also see 39,40]. Cantonese listeners were found to use both pitch contour (i.e., tone shape; falling vs. rising tones) and pitch height (i.e., average height; higher vs. lower tones) to distinguish their native tonal contrasts [35,38]. For example, some Cantonese tone pairs, for instance, T3 (mid-level)-T6 (low-level), are distinguished in subtle differences in pitch height, and thus pose a great perceptual difficulty for listeners [41–43]. In contrast, Mandarin listeners are more sensitive to pitch contour than pitch height [44–47]. Specifically, Mandarin listeners had a great difficulty distinguishing (e.g., Cantonese) level-level tonal contrasts [35,48–51]. Level-level tonal contrasts are perceptually difficult for Mandarin listeners for several reasons: first, a level-level tonal contrast is a novel contrast for Mandarin listeners given that no such contrasts exist in Mandarin tone inventory (see Fig 1); second, Mandarin listeners showed reduced sensitivity to pitch height differences, compared with non-tone language listeners, due to their tonal categorization contrasting in pitch contour [52,53]. Given the Mandarin listeners’ perceptual difficulty, it remains unclear whether a facilitating effect of overnight consolidation in the abstraction would be found for Mandarin listeners’ perceptual learning of Cantonese level-level tonal contrasts, that is, T1 /55/, T3 /33/, and T6 /22/ [33].

Therefore, the primary objective of the present project is to investigate whether overnight consolidation facilitates generalization across talkers, implying the abstraction of novel tonal categories, in the perceptual learning of Cantonese level-level tonal contrasts by Mandarin listeners. Given previous findings on the effect of overnight consolidation in perceptual learning of novel segmental contrasts and the variable/dynamic nature of lexical tones, it is hypothesized that at least a similar facilitating effect of overnight consolidation will be found for perceptual learning of Cantonese level tones by Mandarin listeners. Specifically, those listeners who are trained in the evening are expected to perform better than those who are trained in the morning in perceiving the level tones produced by the new talker and probably also trained talkers, given a potential greater perceptual difficulty of target (three-way) tonal contrasts relative to (two-way) segmental contrasts [54–56]. We also aimed to test whether the overnight consolidation effects found for the identification alone in previous studies [e.g., 16] can be transferred or generalized to the discrimination of tonal contrasts by controlling stimuli variability in both tasks. It is hypothesized that a similar effect will be found in both the identification and discrimination tasks if different degrees of stimuli variability accounted for the discrepancy of the two tasks in terms of the overnight consolidation effect in previous studies. In addition, different from previous studies using a two-way segmental contrast, a three-way tonal contrast is used as our target structure of perceptual learning. Although we do not have specific hypotheses of the overnight consolidation effect with respect to specific level tones or level-level tonal contrasts, it is expected that the effect would be across-the-board for the perceptual learning of all three level tones.

## 2. Materials and Methods

### 2.1 Participants

Thirty-three students (19 female, 14 male) between the ages of 18 and 30 were recruited using an online self-report questionnaire from the Hong Kong Polytechnic University (PolyU). Mandarin-speaking participants who had a minimal exposure to Cantonese were selected based on the following criteria: (1) resided in Hong Kong for less than ten months and learned Cantonese in classroom less than one month prior to the pretest session, (2) speak Mandarin as mother tongue and did not speak any Southern dialect including Shanghainese, Hakka, and Southern Min, and (3) did not receive professional musical training. None of them reported history of hearing impairment, neurological illness, and sleep disorder. One participant was excluded on the basis of a late reporting of exposure to musical (i.e., piano) instruction.

The remaining 32 participants were equally and randomly assigned into one of the two groups who were trained either on the morning (8-10 am) or evening (8-10 pm). Demographic characteristics of the morning and evening groups are summarized in Table 1. A set of pretests was conducted for each group to make sure that the morning and evening groups did not differ at their pitch-related sensitivity and memory capacity at the group level before the training session. The pretest session consists of three tasks: 1). A pitch threshold test adopted from Ho et al. [57] was used to measure the participants’ low-level pitch sensitivities of speech and non-speech tones; 2). A pitch memory span test adopted from Williamson and Stewart [58] was used to measure the participants’ short-term pitch memory capacities; 3). The Montreal Battery of Evaluation of Amusia (MBEA) [59] was used to measure the participants’ musical abilities. The MBEA consists of six subtests: three of them are pitch-based tests (scale, contour, and interval); two of them are duration-based tests (rhythm and meter); the last one is a long-term (musical) memory test. As can be seen in Table 1, the two training groups performed similarly in the pitch threshold task (both speech and non-speech tones), the pitch (short-term) memory span task, the (long-term) memory MBEA test, the pitch-based MBEA tests, and the overall MBEA performance. Results of independent-samples *t*-test confirmed that the scores of the two training groups in each pretest were not significantly different from each other.

**Table 1.** Demographic characteristics of the morning and evening training groups.

|  | **Morning** | **Evening** | ***p*-value** |
| --- | --- | --- | --- |
| No. of participants | 16 (9 F, 7 M) | 16 (9 F, 7 M) | / |
| Age (year) | 24.9 (3.7) | 23.3 (3.5) | *p* =.23 |
| **Pretests** |  |  |  |
| Pitch threshold_speech<br>(in semitone) | 1.6 (2.5) | 1.8 (2.9) | *p* =.86 |
| Pitch threshold_non-speech<br>(in semitone) | 2.4 (3.6) | 1.9 (3.3) | *p* =.66 |
| Pitch short-term memory span<br>(in number of sounds) | 6.2 (1.4) | 6.7 (1.8) | *p* =.34 |
| MBEA_long-term memory (in %) | 91.4 (6.4) | 89.8 (12.4) | *p* =.66 |
| MBEA_pitch-based (in %) | 82.2 (9.8) | 81.1 (9.2) | *p* =.75 |
| MBEA_overall (in %) | 82.7 (8.9) | 81.6 (9.7) | *p* =.82 |

*Note.* The average scores, with the standard deviation in parenthesis, of the two training groups in each pretest are reported. The results ( $p$ -value) of independent-samples $t$ -tests comparing the two groups in age and scores are also reported.

The experimental procedures were approved by the Human Subjects Ethics Sub-committee of the PolyU. Informed written consent was obtained from the participants in compliance with the experiment protocols. All the participants were recruited from January to May 2019, and they were paid for their participation.

### 2.2 Stimuli

The stimuli were 30 words contrasting three Cantonese level tones, /55/ Tone1 (T1, a high-level tone), /33/ T3 (a mid-level tone), and /22/ T6 (a low-level tone). Each tone was carried by ten base syllables (/jan/, /ji/, /jau/, /jiu/, /fan/, /fu/, /ngaa/, /si/, /se/ and /wai/), and all are meaningful in Cantonese. Each monosyllabic target word was embedded in a carrier phrase context “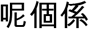_lei1 go3 hai6 [target word]” (this is [target word]). Two female native Cantonese speakers (see Fig 2) recorded the stimuli: tonal stimuli produced by Talker 1 (i.e., the trained talker) were used in the training session; tonal stimuli produced by both Talker 1 and Talker 2 (i.e., the untrained talker) were used in the assessment posttests. Each speaker recorded three repetitions of each target word in the carrier phrase. Recordings were conducted in a sound-proof room using a microphone linked to a digital recorder. Two tokens for each target word were chosen by the investigators based on its intelligibility and sound quality. Each token was segmented out of the carrier phrase, normalized to 500 ms in duration, and scaled to 70 dB SPL in mean intensity using Praat.

**Fig 2.**
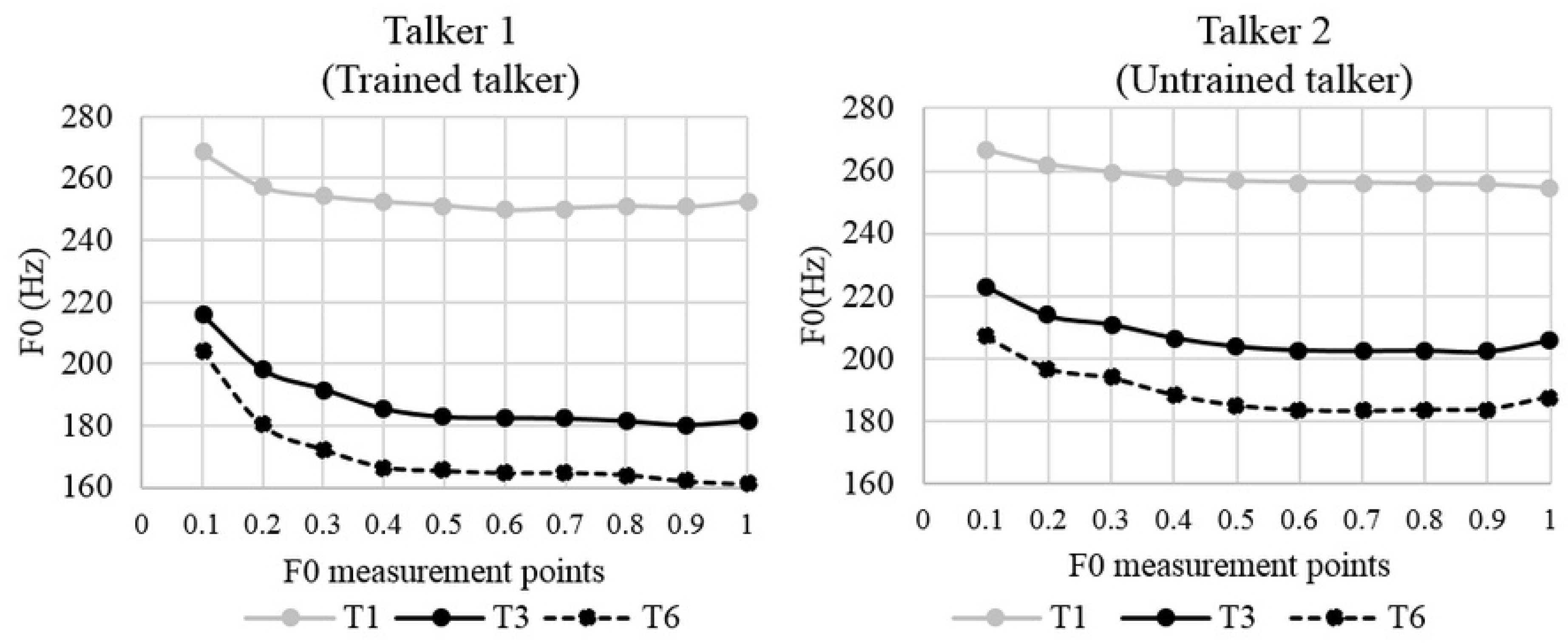
Tonal contours of the three Cantonese level tones. Tonal contours were measured using ten measurement points and produced by the trained female talker (left panel) whose stimuli were used in both the training and posttests as well as the untrained female talker (right panel) whose stimuli were used in the posttests alone.

The untrained talker differed phonetically from the trained talker in their voice similarity and pitch distribution, and thus was used as a generalization talker in this study. The trained and untrained talkers were selected from six female talkers used in another project [60]. First, the untrained talker, along with other four female talkers, was compared against the trained talker in terms of voice similarity. The similarity was rated by 12 native Cantonese speakers on a scale of 1 to 9 (with 1 meaning very dissimilar and 9 meaning very similar). The untrained talker received the lowest similarity rating score (i.e., 4.70) averaged across raters among five talkers (rating range: 4.70-7.44 on a scale of 1-9) when rated against the trained talker. In addition, as illustrated in Fig 2, the trained and untrained talkers had different pitch distribution with the latter having a higher average pitch (trained: 204 Hz; untrained: 219 Hz) and a narrower pitch range (trained: 170-254 Hz; untrained: 189-258 Hz). Due to the between-talker F0 difference, T6 (low-level tone) produced by the untrained talker had even higher pitch values than T3 (midlevel tone) produced by the trained talker. Therefore, given the voice and pitch distribution dissimilarity between the two talkers, generalizing across the trained and untrained talkers was essential for trainees to learn how to use pitch height in categorizing novel level tones after perceptual training.

### 2.3 Procedures

Following the design of previous studies [16,20], a pretest-training-posttest paradigm was conducted on the participants in the two training groups, a morning training group vs. an evening training group. As shown in Table 2, the morning group was trained in the morning (8-10 am) in Session 1, and then tested with a 12-hours delay (8-10 pm) in Session 2 over the course of one day. In contrast, the evening training group was trained in the evening (8-10 pm) in Session 1, and then tested with an intervening night’s sleep after a similar time delay (8-10 am) in Session 2 across two consecutive days. Participants completed a sleep questionnaire for the 24-hour experiment period at the end.

**Table 2.** Overview of timing in the experimental protocol.

| *Days* | Day1 | Day 2 |  | Day 3 |  |
| --- | --- | --- | --- | --- | --- |
| *Time* |  | **8-10AM** | **8-10PM** | **8-10AM** | **8-10PM** |
| *Sessions* | **Pretest session** | **Session1** | **Session2** | **Session3** |  |
| **Morning Training Group** | 1.Pitch threshold task | AX pretest |  |  |  |
|  | 2.Pitch memory task | ID training |  |  |  |
|  | 3. MBEA | ID posttest1<br>(immediate) | ID posttest2<br>(12-hour delay) | ID posttest3<br>(24-hour delay) |  |
|  |  | AX posttest1<br>(immediate) | AX posttest2<br>(12-hour delay) | AX posttest3<br>(24-hour delay) |  |
| **Evening Training Group** | 1.Pitch threshold task |  | AX pretest |  |  |
|  | 2.Pitch memory task |  | ID training |  |  |
|  | 3. MBEA |  | ID posttest1<br>(immediate) | ID posttest2<br>(12-hour delay) | ID posttest3<br>(24-hour delay) |
|  |  |  | AX posttest1<br>(immediate) | AX posttest2<br>(12-hour delay) | AX posttest3<br>(24-hour delay) |

The experiments were conducted using Paradigm software (Perception Research Systems, Inc. http://www.paradigmexperiments.com/). Auditory stimuli were presented to the participants via stereo headphones with the volume adjusted to a comfortable level. The participants completed the experiments in a quiet space in the Speech and Language Sciences Lab of PolyU.

As mentioned in 2.1, a set of pretests was conducted for each group before training on a separate day (i.e., Day 1). In the training session, a self-paced, forced-choice identification (ID) task of Cantonese level tonal contrasts was conducted on the two training groups. Monosyllables produced by the trained talker (Talker 1 in Fig 2) alone were used in the training session. During the training, the participants were instructed to identify each tone (T1-High, T3-Mid and T6-Low) by pressing three buttons (1, 3, and 6). A total of 300 trials (1 talker * 3 tones * 10 syllables * 2 tokens * 5 repetitions) were auditorily presented to the participants in five blocks with 60 trials in each block. Written feedback (“Correct” in green or “Incorrect. The correct answer is…” in red) was given immediately after every trial. The participants were instructed to learn to categorize three tones based on feedback, and achieve the best performance as they can in this session.

In the posttest sessions, the participants were assessed on both ID and AX discrimination. As shown in Table 2, each posttest was conducted at three time points: Immediately after training, with a 12-hour delay, and with a 24-hour delay, to test how the learned Cantonese level tones were retained and how the mental representation of learned tones changed over time after perceptual training. Monosyllables produced by both the trained (Talker 1 in Fig 2) and untrained (Talker 2 in Fig 2) talkers were used in the posttest sessions.

In the ID posttests, the participants were instructed to identify each tone (T1-High, T3-Mid and T6-Low) by pressing three buttons (1, 3, and 6). However, no feedback was given after every trial in the posttests. The ID posttest consisted of 120 trials (2 talkers * 3 tones * 10 syllables * 2 tokens), which were randomly presented to participants in one block.

Following each ID posttest, participants were tested in an AX discrimination posttest. In the AX discrimination posttests, the participants were instructed to distinguish whether two tones they heard belong to the same or different tone categories by pressing one of two buttons (left arrow and right arrow) indicating “same” or “different”, respectively, on the keyboard. No feedback was given in the posttests. An equal number of AA pairs (120 pairs with the same tone within each pair) and AB pairs (120 pairs with different tones within each pair) were used to counterbalance the two types of tone pairs. The presentation order of two tones in each AB pair was also counter-balanced in different trials. Two acoustically different tokens of the same tone type were used in each AA pair. In addition, a relatively long inter-stimulus interval, 1000 ms, was used. The design of the AX posttest allowed us to tap an individual’s recognition of the tone category rather than the use of low-level acoustic information (e.g., pitch) to discriminate tonal tokens. The AX posttest had 240 trials (2 talkers *3 pairs * 2 orders * 10 syllables * 2 types), which were presented in a random order to the participants in one block.

Following the design of [16], an AX discrimination pretest was conducted as a baseline assessment to make sure that discrimination ability of the two training groups is comparable before the training session. In total, the study involves three sessions of behavioral experiments requiring four lab visits for each participant, which took about four hours in total.

### 2.4 Data analysis

For the ID task, a logit mixed-effects model with Group (2 levels: morning vs. evening; morning as Baseline), Time (3 levels: ID posttest1, ID posttest2, ID posttest3; ID posttest1 as Baseline), Talker (2 levels: trained vs. untrained; trained talker as Baseline), and Tone (3 levels: T1, T3, and T6; T1 as Baseline) as fixed effects, with participants and items as random intercepts and Time as a random slope on the participant random intercept (the models with more random slopes did not converge), was performed on participants’ accuracy data (1 = correct, 0 = incorrect) of their ID posttests.

For the AX discrimination task, a linear mixed-effects model with Group (2 levels: morning vs. evening; morning as Baseline), Time (4 levels: pretest, posttest1, posttest2, posttest3; pretest as Baseline), Talker (2 levels: trained vs. untrained; trained talker as Baseline), and Tone Pair (3 levels: T1-T3, T1-T6, and T3-T6; T1-T3 as Baseline) as fixed effects, with participants (item information was obscured by computing *d*-prime scores of the discrimination task, so item was not in the random effect structure) as a random intercept and Time as a random slope (the models with more random slopes did not converge), was conducted on participants’ *d*-prime scores of their AX discrimination posttests. A *d*-prime score was used to assess the participants’ discrimination ability. The *d*-prime score for each participant was derived based on the “hit” rate (number of times the “different” button was pressed for AB pairs) and the “false alarm” rate (number of times the “different” button was pressed for AA pairs). *D*-prime score is the difference between “hit” rate and the “false alarm” rate when they are *z*-transformed [61].

The models were fitted in R, using the glmer() and lmer() function from the lme4 package for logit and linear mixed-effects models, respectively [for discussion, 62]. A back-fitting function from the package *LMERConvenienceFunctions* in R [63] was used to identify the best model that accounted for significantly more of the variance than simpler models, as determined by log-likelihood ratio tests; only the results of the model with the best fit are presented, with *p* values calculated using the *lmerTest* package in R [64]. Analyses yielding significant interactions between fixed factors (e.g., Group and Time) were followed up by subsequent models conducted separately, for instance, for each group.

If there are sleep-mediated changes in the identification and discrimination maintenance (i.e., the overnight consolidation effect), the morning and evening training groups will differ in performance changes over the 24-hour experiment period, yielding a significant two-way interaction between Group and Time in both ID and AX discrimination tasks. If the potential sleep-mediated changes were only found for stimuli produced by the untrained talker, the performance changes of the morning and evening training groups over time will differ between stimuli produced by the trained and untrained talker, yielding a significant three-way interaction of Group, Time and Talker in both ID and AX discrimination tasks. We do not predict that the overnight consolidation differs across specific level tonal contrasts, so the potential interaction between Group and Time might not interact with Tone in the ID tasks and Tone Pair in the AX discrimination tasks.

## 3. Results

To ensure that the morning and evening groups had comparable levels of tone discrimination sensitivity before going through training, an independent-samples *t*-test by Group was performed on the accuracy of the AX discrimination pretest, a baseline assessment. The accuracy of the morning and evening groups did not significantly differ from each other [*t*(30) = −.46, *p* = .65].

We do not have a baseline measure of ID performance, because the tone-category pairing would have been random before the participants learned to categorize tones in the training session. To ensure that participants performed above chance following training, a one-sample *t*-test was conducted on the participants’ accuracy in the ID posttest1 immediately after training. The accuracy of the morning group [*t*(15) = 8.80, *p* < .001] and the evening group [*t*(15) = 9.72, *p* = < .001] in the ID Posttest were both statistically above chance, 33.3%. Moreover, the results of an independent-samples *t*-test by Group on the accuracy of the ID posttest1 showed that differences in group performance immediately following training were not statistically significant [*t*(30) = −.61, *p* = .55], implicating that the degree of potential improvement immediately after training was comparable between the morning and evening groups.

### 3.1 ID Performance

Fig 3 presents the morning and evening training groups’ mean proportions of correct response (i.e., accuracy) for stimuli produced by the trained and untrained talkers over the 24-hour period (ID posttest1, ID posttest2, and ID posttest3) in the ID posttests.

**Fig 3.**
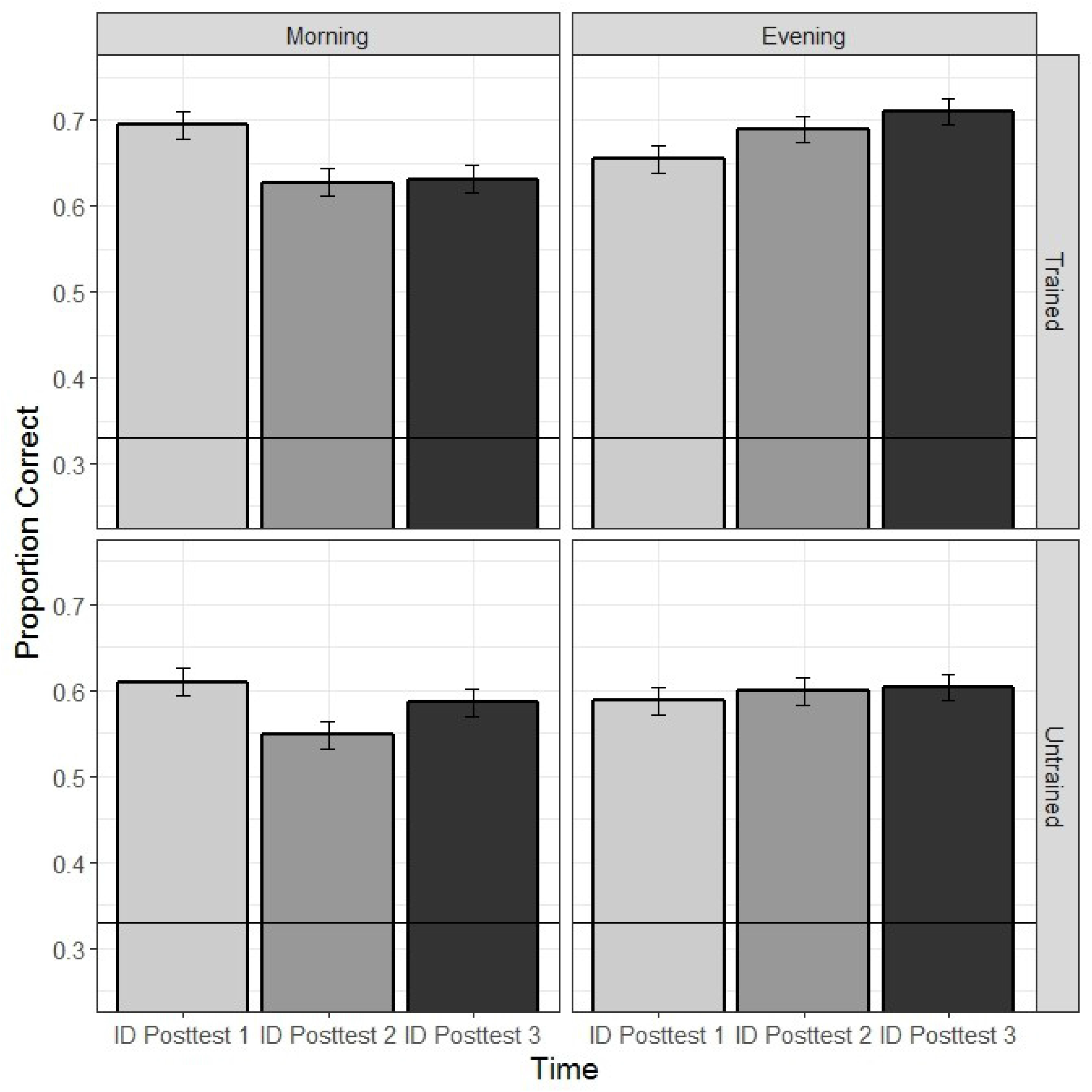
The morning and evening training groups’ mean proportions of correct response for stimuli produced by the trained and untrained talkers over the 24-hour period in the ID posttests. The error bars represent one standard error of the mean; the horizontal line represents chance performance.

Recall that a logit mixed-effects model was performed on participants’ accuracy to examine the effects of Group (morning vs. evening), Time (ID posttest1, ID posttest2, ID posttest3), Talker (trained vs. untrained), and Tone (T1, T3, and T6), and their interactions. The baseline was the morning groups’ performance on trained stimuli of T1 in the ID posttest1. The model with the best fit included the simple effects of Group, Time, Tone, Talker, as well as the interaction between Group and Time. The estimate, standard error, *z* value, and *p* value associated with the fixed effects are presented in Table 3.

**Table 3.**
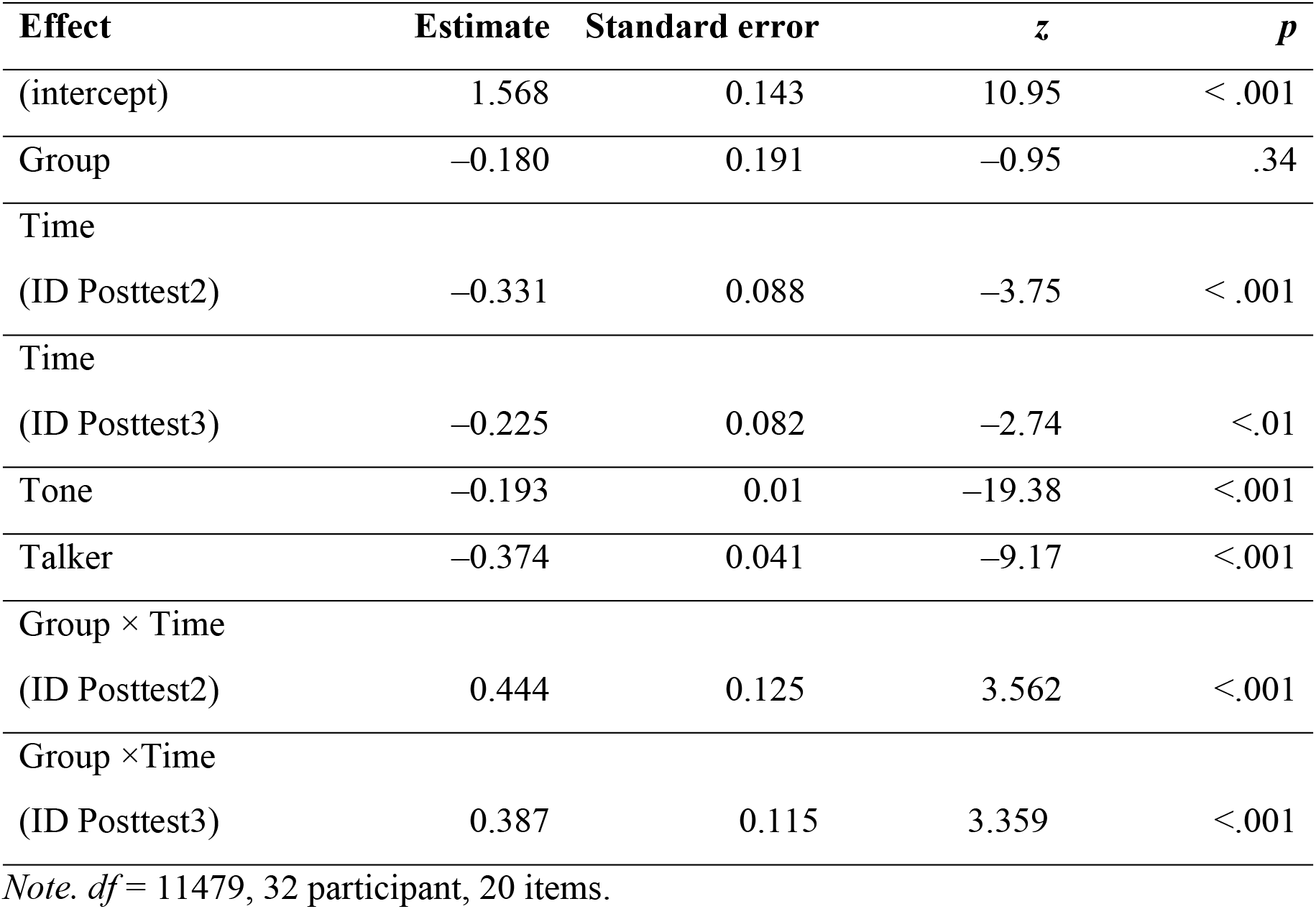
Best logit mixed-effects model on proportions of correct response of the participants from the morning and evening groups in the three AX posttests.

The model results summarized in Table 3 indicate that the morning groups’ ID performance on trained stimuli of T1 in the ID posttest1 is larger than 0 (intercept); their performance (of accuracy) in either ID posttest2 or 3 was lower than their performance in the ID posttest1; their performance of T3 and T6 was lower than their performance of T1 trials; their performance for the untrained talker was lower than their performance for the trained talker. Importantly, the model also yielded a significant two-way interaction between Group and Time, indicating that the morning and evening training groups differed in their ID performance changes over the 24-hour experiment period. However, the three-way interaction among Group, Time and Talker did not improve the model, indicating a lack of evidence that the ID performance changes over time of the morning and evening training groups differed between stimuli produced by the trained and untrained talkers.

To understand the nature of the significant two-way interaction between Group and Time, subsequent logit mixed-effects models were therefore performed on the participants’ accuracy across three ID posttests separately for each training group. For the morning group, the participants’ performance in either the ID posttest2 (Estimate = −0.321, Std. Error = 0.069, *z* = −4.65, *p* < 0.001) or the ID posttest3 (Estimate = −0.224, Std. Error = 0.070, *z* = −3.22, *p* <. 01) was significantly lower than their performance in the ID posttest1. However, the participants’ performance between the ID posttest2 and the ID posttest3 did not significantly differ (Estimate = 0.100, Std. Error = 0.068, *z* = 1.47, *p* = 0.14). The results of the morning group suggest that participants without an intervening night’s sleep between Session 1 (i.e., training session) and Session 2 (i.e., posttest2) showed a declining ID performance in the later posttests compared with their ID performance in the initial posttest.

For the evening group, the participants’ performance in the ID posttest2 did not differ from their performance in the ID posttest1 (Estimate = 0.100, Std. Error = 0.069, *z* = 1.47, *p* =. 14). Importantly, the participants’ performance in the ID posttest3 was significantly higher than their performance in the ID posttest1 (Estimate = 0.147, Std. Error = 0.069, *z* = 2.13, *p* =. 03). Again, the participants’ performance in the ID posttest2 and the ID posttest3 did not significantly differ (Estimate = 0.046, Std. Error = 0.070, *z* = 0.66, *p* = 0.51). The results of the evening group suggest that participants with an intervening night’s sleep between Session 1 (i.e., training session) and later sessions showed a trend of improved ID performance in posttest2, and significant improvement in posttest3, compared with their ID performance in the initial posttest.

### 3.2 AX Discrimination Performance

Fig 4 presents the morning and evening training groups’ mean *d*-prime scores for stimuli produced by the trained and untrained talkers over the 24-hour period (AX pretest, AX posttest1, AX posttest2, and AX posttest3) in the AX discrimination.

**Fig 4.**
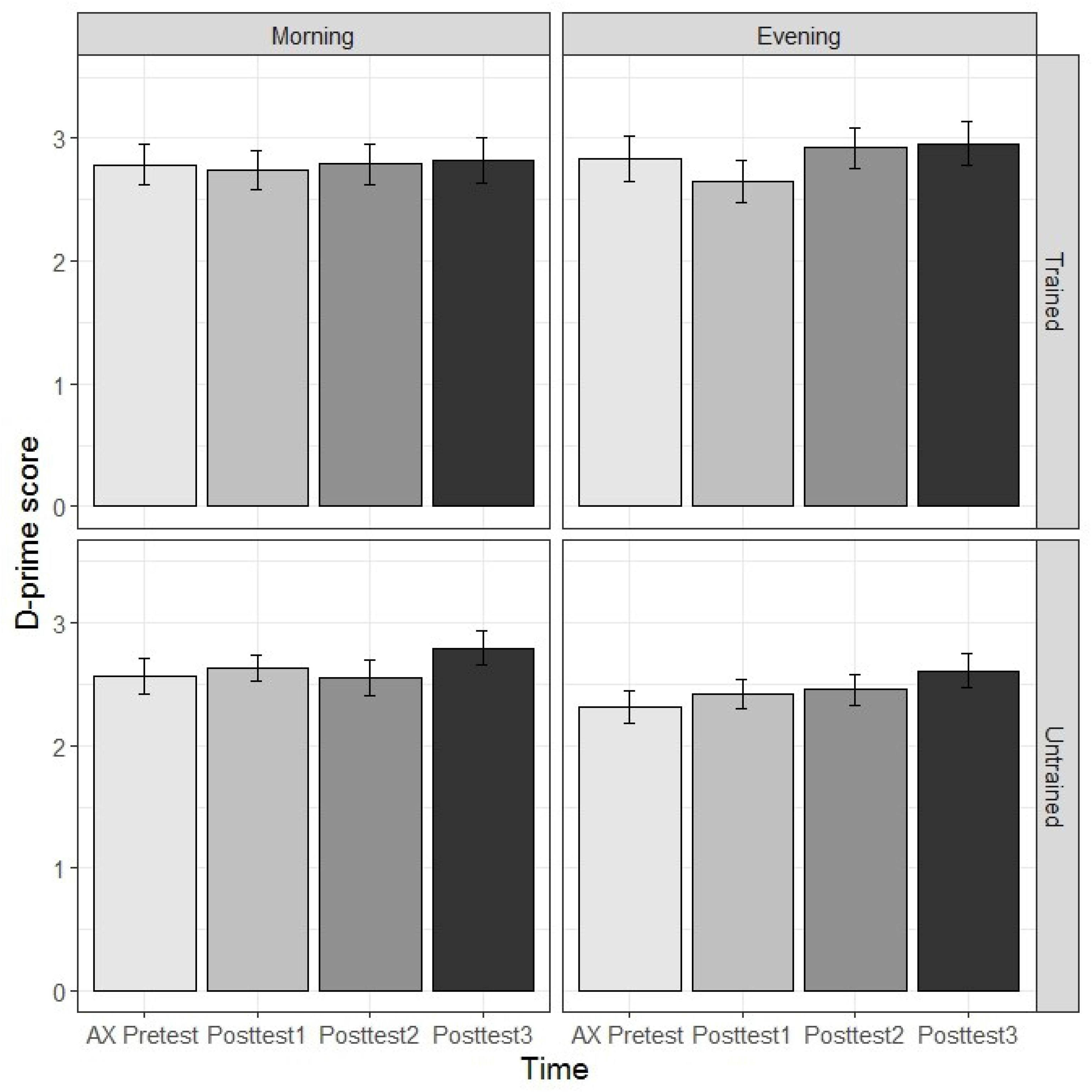
The morning and evening training groups’ mean *d*-prime scores for stimuli produced by the trained and untrained talkers over the 24-hour period in the AX discrimination pretest and posttests. The error bars represent one standard error of the mean

Recall that a linear mixed-effects model was performed on participants’ *d*-prime scores to examine the effects of Group (morning vs. evening), Time (AX pretest, AX posttest1, AX posttest2, and AX posttest3), Talker (trained vs. untrained), and Tone Pair (T1-T3, T1-T6, and T3-T6), and their interactions. The baseline was the morning groups’ performance on trained stimuli of T1-T3 in the AX discrimination pretest. The model with the best fit included the simple effects of Group, Tone Pair, Talker, and the interaction between Group and Talker as well as between Tone Pair and Talker. The estimate, standard error, *t* value, and *p* value associated with the fixed effects are presented in Table 4.

**Table 4.** Best linear mixed-effects model on *d*-prime scores of the participants from the morning and evening groups in the pretest and three posttests of AX discrimination.

| Effect | Estimate | Standard error | *df* | *t* | *p* |
| --- | --- | --- | --- | --- | --- |
| (intercept) | 3.245 | 0.121 | 44.86 | 26.76 | < .001 |
| Group | 0.056 | 0.162 | 36.06 | 0.34 | .73 |
| Tone Pair |  |  |  |  |  |
| (T1-T6) | 0.351 | 0.068 | 736 | 5.18 | <.001 |
| Tone Pair |  |  |  |  |  |
| (T3-T6) | -1.729 | 0.068 | 736 | -25.53 | <.001 |
| Talker | -0.408 | 0.078 | 736 | -5.22 | <.001 |
| Group × Talker | -0.243 | 0.078 | 736 | -3.10 | <.01 |
| Tone Pair × Talker |  |  |  |  |  |
| (T1-T6) | 0.143 | 0.096 | 736 | 1.49 | .14 |
| Tone Pair × Talker |  |  |  |  |  |
| (T3-T6) | 0.634 | 0.096 | 736 | 6.62 | <.001 |

The model results summarized in Table 4 indicate that the morning group’s performance on trained stimuli of T1-T3 trials in the AX discrimination task is larger than 0 (intercept); their performance of T1-T6 pairs was higher than their performance of T1-T3 pairs; their performance of T3-T6 pairs was lower than their performance of T1-T3 trials; their performance for the untrained stimuli was lower than their performance for the trained stimuli.

The model also yielded significant two-way interaction effects of Group and Talker as well as of Tone Pair (T3-T6) and Talker. Crucially, the interaction between Group and Time did not improve the model, indicating that the morning and evening training groups did not differ in AX discrimination performance changes over the 24-hour experiment period. The three-way interaction among Group, Time and Talker did not improve the model neither, indicating that the morning and evening training groups showed similar discrimination performance changes over time for stimuli produced by both the trained and untrained talkers.

To understand the nature of the significant two-way interaction between Group and Talker, subsequent linear mixed-effects models were therefore performed on the participants’ *d*-prime scores for stimuli produced by the trained and untrained talkers separately for each training group. As illustrated in Fig 4, for the morning group, the participants’ performance for the trained and untrained talkers did not significantly differ (Estimate = −0.150, Std. Error = 0.101, *t* = −1.48, *p* = 0.14). For the evening group, the participants’ performance for the trained talker was significantly higher than their performance for the untrained talker (Estimate = −0.392, Std. Error = 0.098, *t* = −3.99, *p* < 0.001). The results suggest that the evening group had greater difficulty discriminating the stimuli produced by the untrained talker, who had a narrower pitch range as well as smaller pitch differences between some tones (e.g., T1 and T3) than the trained talker (see Fig 2). Importantly, the perceptual pattern was the same across the pretest and posttests without Time modulating it. Thus, the finding was not attributed to an overnight consolidation effect in terms of the talker generalization/abstraction.

To understand the nature of the significant two-way interaction between Tone Pair (T3-T6) and Talker, subsequent linear mixed-effects models were therefore performed on the participants’ *d*-prime scores for stimuli produced by the trained and untrained talkers separately for T1-T3 and T3-T6. As illustrated in Fig 5, for T1-T3 trials, the participants’ performance for stimuli produced by the untrained talker was significantly lower than their performance for stimuli produced by the trained talker (Estimate = −0.530, Std. Error = 0611, *t* = −8.66, *p* < 0.001). For T3-T6 trials, the participants’ performance for stimuli produced by the trained and untrained talker did not significantly differ (Estimate = 0.104, Std. Error = 0.056, *t* = 1.87, *p* = .06). The results suggest that all the participants found T1-T3 trials produced by the untrained talker, which had smaller pitch differences than T1-T3 trials produced by the trained talker (see Fig 2), perceptually more difficult than the counterparts of the trained talker. Importantly, the perceptual pattern was the same across the training groups and the pretest-posttests without Group (groups with vs. without intervening night’s sleep between training and posttest2) and Time modulating it. Thus, the finding was not attributed to an overnight consolidation effect.

**Fig 5.**
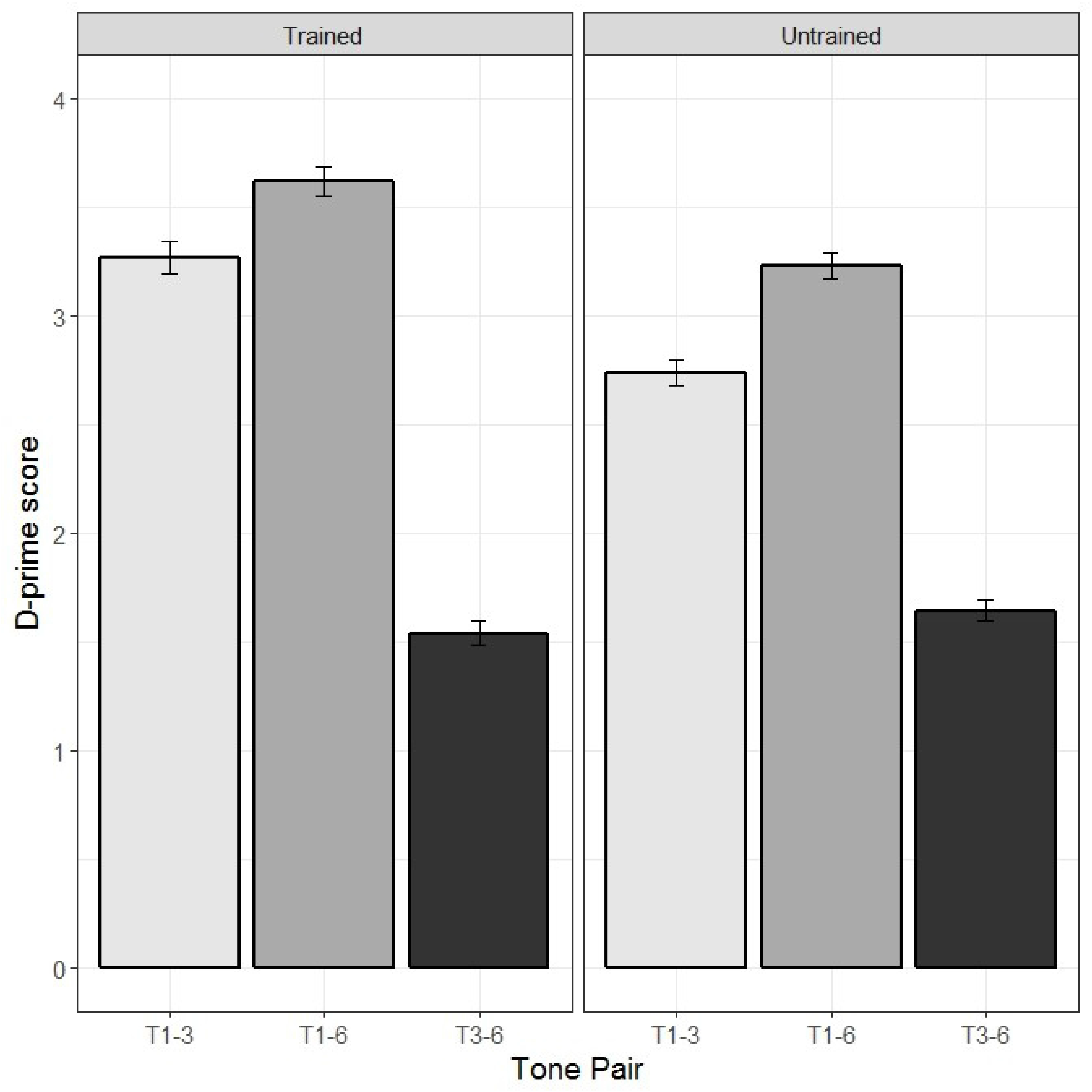
The participants’ mean *d*-prime scores of T1-T3, T1-T6, and T3-T6 trials produced by the trained talker (left panel) and the untrained talker (right panel). The error bars represent one standard error of the mean.

In summary, the morning and evening training groups differed in their ID performance changes over the 24-hour experiment period. More specifically, participants in the morning group, without an intervening night’s sleep between training and ID posttest2, showed a declining ID performance of accuracy in the later posttests compared with their ID performance in the initial posttest. In contrast, the participants in the evening group, having an intervening night’s sleep between training and later sessions, showed a trend of improved ID performance of accuracy in ID posttest2, and significant improvement in ID posttest3 compared with their ID performance in the initial posttest. Importantly, the different perceptual patterns over time between the morning and evening training groups were found for the stimuli produced by both the trained and untrained talkers. Different from the ID performance, the morning and evening training groups did not show divergent performance changes of the AX discrimination tasks over time. The evening group’ lower discrimination performances (in *d*-prime scores) of stimuli produced by the untrained talkers than those by the trained talker, as well as all the participants’ lower discrimination performance performances (in *d*-prime scores) of T1-T3 trials produced by the untrained talker than those by the trained talker are likely to be driven by the acoustic difference of tonal stimuli produced by the trained versus the untrained talkers as well as their less familiarity with the untrained talker.

### 3.3 The Effect of Sleep-related Variables

This analysis focused on the evening group to examine the relation of their performance changes over the 24-hour period to their sleep. The results of the sleep questionnaire showed that the participants in the evening group had a sleep duration at an average of 7.3-hour (range: 6-8 hours) between session 1 (training) and session 2 (posttest2). They got an average of 7.6 (range: 6-9) as a self-rating score of their sleep quality on a 1-10 point scale (with 1 meaning very bad and 10 meaning very good). The participants self-reported to fall asleep at 00:40 am on average (range: 11:30 pm to 2:30 am) in the evening, and to wake up at 7:50 am on average (range: 7:00 am - 8:30 am) in the morning.

A set of regression analyses were carried out to examine to what extent the performance changes of the participants in the evening group can be predicted by their sleep-related variables. Because sleep-related variables are highly collinear, self-reported sleep duration, sleep quality rating, sleep time (what time did the participants fall asleep) and wake-up time (what time did the participants wake up) were entered stepwise as predictors into a multiple regression model using SPSS software with performance changes (the difference between posttest2 and posttest1; the difference between posttest3 and posttest1) as the dependent variables for each task.

The results of regression analyses showed, as illustrated in Fig 6, that sleep time accounted for the largest proportion of variance in overnight identification changes, that is, the difference between posttest2 and posttest1 [F (1.14) = 5.991, *p* = .028, r^2^ = .537]. Sleep time also accounted for the largest proportion of variance in the identification changes between posttest3 and posttest1 [F (1.14) = 6.992, *p* = .019, r^2^ = .577]. No other sleep-related variables significantly accounted for the variance in discrimination changes. The findings suggest that the earlier the participants fell asleep, the more identification gains they would have in the ID posttests on the following day.

**Fig 6.**
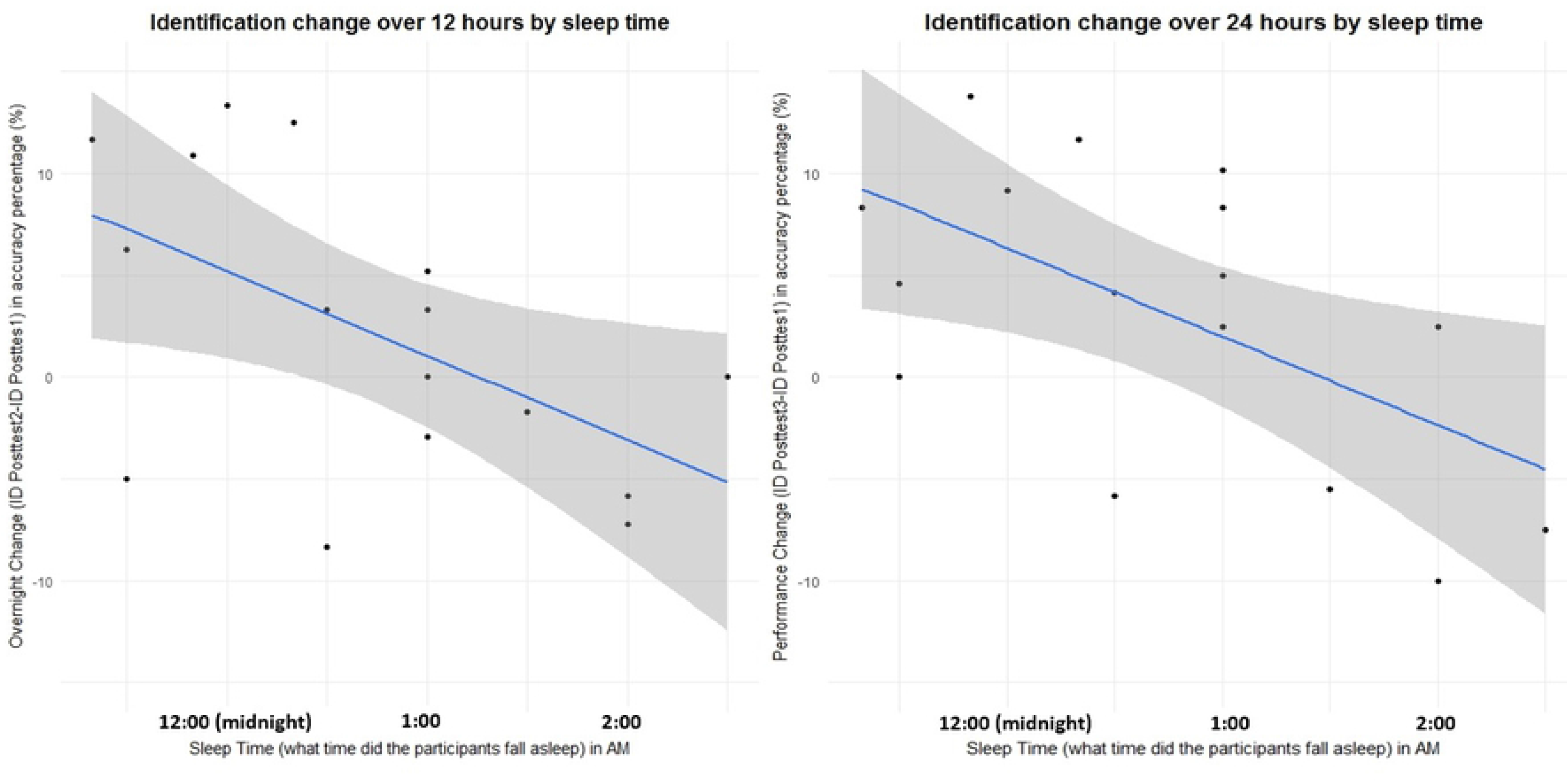
Relationships between individual differences in sleep time (in AM) and the performance (accuracy percentage in %) changes over 12 hours (left) and 24 hours (right) to the identification task performance. The shaded areas represent a 5% confidence interval.

## 4. Discussion and conclusion

The current study examined whether overnight consolidation facilitates generalization across talkers, implying an abstraction of novel tonal information, in the identification and discrimination of novel Cantonese level-level tonal contrasts for Mandarin listeners. The results of identification tasks showed that the Mandarin listeners who were trained in the morning, without an intervening night’s sleep between training and later posttests, showed a declining performance of identification accuracy; in contrast, the Mandarin listeners who were trained in the evening, with an intervening night’s sleep between training and later posttests, showed improved performance in identification accuracy. Crucially, the pattern was found for stimuli produced by both the trained and untrained talker across all three level tones. Moreover, the identification performance changes of the participants in the evening group were predicted by how early they fell asleep right after the training session. On the other hand, the results of discrimination tasks did not reveal divergent performance changes for the two training groups.

First and foremost, the findings suggest that overnight consolidation facilitates generalization across talkers in the identification of novel Cantonese level-level tonal contrasts by Mandarin listeners. More specifically, sleep-mediated overnight consolidation might have assisted the evening trainees’ generalization of acoustic-phonetic (i.e., pitch) features from the stimuli produced by the trained talker to those produced by the untrained talker, and eventually facilitated their formation of a more abstract representation of Cantonese level tone categories in memory traces (e.g., in the hippocampus of human brains). This account is in line with the findings of previous research showing that overnight consolidation promoted listeners’ talker generalization or abstraction for perceptual learning and lexically-guided phonetic retuning of non-native segments [14,16,20].

In addition, the significant relationship between the identification performance changes of the evening trainees and their sleep time provided further evidence supporting the account of overnight consolidation. The findings suggest that the earlier the trainees fell asleep, the more likely it was for a facilitating effect of overnight consolidation to emerge in the perceptual learning of Cantonese level tones. This finding is also consistent with that of recent studies showing that sleep-related variables (e.g., total sleep duration) predicted behavioral and neural changes of adult speech learning of non-native segments after a perceptual training in the evening [17]. While an earlier sleep time is an important indicator of and possibly correlated with a higher sleep quality and/or a longer sleep duration, self-rated sleep quality scores and selfreported sleep duration were not found to predict the identification performance changes of the evening trainees in the current study. It is possible that the questionnaire method adopted in the current study renders the sleep duration and quality data less precise, and an objective tool (e.g., a sleep-monitoring headband) is thus suggested to be used to further investigate whether, and if so how, different sleep-related variables affect the evening trainees’ performance changes over time. Future studies may also recruit a larger group of participants with a broader range of variability in sleep duration and quality.

Another possibility to account for the divergent performance changes of the morning and evening groups could be attributed to the presence of interference from Mandarin tone input (or any other speech input) or not outside the laboratory after a perceptual training [16,19,20]. According to this account, while the morning group’s exposure to Mandarin tones (and other speech input) subsequent to training interfered with the consolidation of Cantonese level tones and resulted in a decline of identification performance, the evening group’s overnight sleep interval prevented such interference. While it is necessary to acknowledge that this possibility cannot be ruled out with a design of comparing the morning and evening training groups, our findings did not seem to fully support this account. First, our results showed that the identification performance of the evening group after 24-hour (i.e., in ID posttest3) did not decline, and remained stable instead compared with earlier posttests (e.g., ID posttest2), after an exposure to Mandarin tones (and other speech input) during a daytime interval between posttest2 and posttest3. Second, based on the interference account, the morning group was expected to have a declining performance over time due to the interference effect; in contrast, the evening group, without such interference, was not necessarily expected to show improved performance which can be predicated by sleep-related variables (i.e., how early they fell asleep). Therefore, while the results should be interpreted cautiously given this alternative account, we propose that the effect of overnight consolidation should have, at least partially, contributed to the divergent changes of identification performance by the two training groups in the present study. Future research is suggested to focus on participants who are trained in the evening to get around this possibility in examining the effect of overnight consolidation (e.g., with sleep-related variables of evening trainees monitored) in perceptual learning of non-native sounds.

Importantly, the results showed that the facilitation effect of overnight consolidation was found for stimuli produced by both the trained and untrained talkers. An implication of the results is that the overnight consolidation might have a larger facilitation effect of abstraction in perceptual learning of lexical tones than that of segments, given the variable and dynamic nature of lexical ones. Different from the results of the previous study [16] which found the different perceptual changes of the two training groups in identifying the stop stimuli produced by the untrained talker alone, we found that divergent performance changes of the two groups in identifying the level-tone stimuli produced by both the trained and untrained talkers. The discrepancy between this study and the previous study can be potentially attributed to the different nature of segments (e.g., stop) and lexical tones. One possibility is that the variable and dynamic nature of lexical tones made it more difficult for participants to learn/abstract than segments. As a result, the intervening sleep between training and later posttests thus facilitated the stimuli produced by the trained talker, which were also perceptually challenging [22,23,26,27].

An alternative (but related) possibility to account for the discrepancy regarding the talker effect is that the training stimuli in the current study had a greater variability than those in the previous study [16]. While this study used two tokens of each tone carried by ten different syllables as training stimuli, the previous study [16] used five tokens of each stop carried by a single syllable (i.e., vowel context). In addition, three-way tonal contrasts were trained as target stimuli in the current study whereas a two-way segmental contrast was used in the previous study. Thus, a greater variability of training stimuli in this study than those used in the previous study [16] might have made the novel tonal categories perceptually more challenging, and thus contribute at least partially to the discrepancy regarding the effect for the trained talker’s stimuli. Previous tone training studies found that stimulus variability of syllables and speaker would modulate the perceptual learning outcome [48,65–68]. Thus, a future direction of this line of research is to investigate whether a greater or a smaller acoustic variability (of syllables and/or speakers) of training stimuli would modulate the size of the overnight consolidation effect in facilitating the abstraction of novel phonetic information.

An important question that the current results raise is why the two training groups showed divergent performance changes over time in the identification tasks but not in the discrimination tasks. First, our results are consistent with those in previous studies on segments [16,19], which claimed that the lack of group differences in the discrimination tasks was probably due to its greater stimuli variability (i.e., untrained vowel contexts) than those in the identification tasks. The findings of the current study did not seem to support this account as identical tonal stimuli were used in both the identification and discrimination tasks. Thus, it is possible that the overnight consolidation effect in terms of abstraction is more related to the identification than the discrimination of novel sound categories in nature. One more plausible explanation for the lack of effects in the discrimination tasks is the different natures of training (identification) and assessment (discrimination) tests, which tapped into different aspects of non-native tone perception. The identification task used in the training session (and posttests) tapped into higher levels of phonological encoding of tonal categories. In contrast, the discrimination tasks used in the posttests, which were intended to tap an individual’s phonological processing of tone categories by using two different tokens of the same tone category in AA pairs and a relatively long ISI, might have tapped into relatively low levels of phonetic processing instead. The phonetic processing in sound discrimination did not change much even after multiple perceptual training sessions according to previous training studies [69–71]. A discrimination task with feedback in the training session can be used in further studies to examine whether an effect of overnight consolidation is restricted to high-level phonological abstraction in nature or can be ultimately yielded in the low-level sensitivity assessments (e.g., discrimination).

To the best of our knowledge, the present study is the first to examine the effect of overnight consolidation in perceptual learning of lexical tones. The present findings suggest that sleep-mediated overnight consolidation (at least partially) facilitated the perceptual learning of Cantonese level tones by the Mandarin-speaking evening trainees, in promoting their generalization or abstraction from the trained talker to the untrained talker, and crucially yielding an across-the-board improvement of their identification performance over the 24-hour experiment period. These findings spark interest in several questions regarding training stimuli, sleep-related variables and others in further research. Among other things, it would be intriguing to investigate the potential effect of overnight consolidation in perceptual learning of different types of tone pairs (e.g., contour-contour tone pairs vs. level-level tone pairs) and other prosodic categories (e.g., lexical stress) to further shed lights on the effect of overnight consolidation in perceptual learning of suprasegmental domain as well as different nature of segments and prosodic categories in this process.

## Acknowledgements

This work was supported in part by the Departmental General Research Funds (International collaboration) and the Departmental Reward Scheme for Research Publications in Indexed Journals awarded to CCZ, and the Language Learning Early Career Research Grant and the Postdoctoral Fellowships Scheme at the Department of Chinese and Bilingual Studies of the Hong Kong Polytechnic University awarded to ZQ.

